# CAGECAT: The CompArative GEne Cluster Analysis Toolbox for rapid search and visualisation of homologous gene clusters

**DOI:** 10.1101/2023.02.08.527634

**Authors:** Matthias van den Belt, Cameron Gilchrist, Thomas J. Booth, Yit-Heng Chooi, Marnix H. Medema, Mohammad Alanjary

## Abstract

**Background:** Co-localized sets of genes that encode specialized functions are common across microbial genomes and occur in genomes of larger eukaryotes as well. Important examples include Biosynthetic Gene Clusters (BGCs) that produce specialized metabolites with medicinal, agricultural, and industrial value (e.g. antimicrobials). Comparative analysis of BGCs can aid in the discovery of novel metabolites by highlighting distribution and identifying variants in public genomes. Unfortunately, gene-cluster-level homology detection remains inaccessible, time-consuming and difficult to interpret.

**Results:** The comparative gene cluster analysis toolbox (CAGECAT) is a rapid and user-friendly platform to mitigate difficulties in comparative analysis of whole gene clusters. The software provides homology searches and downstream analyses without the need for command-line or programming expertise. By leveraging remote BLAST databases, which always provide up-to-date results, CAGECAT can yield relevant matches that aid in the comparison, taxonomic distribution, or evolution of an unknown query. The service is extensible and interoperable and implements the cblaster and clinker pipelines to perform homology search, filtering, gene neighbourhood estimation, and dynamic visualisation of resulting variant BGCs. With the visualisation module, publication-quality figures can be customized directly from a web-browser, which greatly accelerates their interpretation via informative overlays to identify conserved genes in a BGC query.

**Conclusion:** Overall, CAGECAT is an extensible software that can be interfaced via a standard web-browser for whole region homology searches and comparison on continually updated genomes from NCBI. The public web server and installable docker image are open source and freely available without registration at: https://cagecat.bioinformatics.nl

## BACKGROUND

Genes working cooperatively in a metabolic pathway are often physically co-localized in prokaryotic and fungal genomes. These gene clusters are commonly observed in specialized metabolism involved in ecological adaptations, such as nutrient utilization and production of virulence factors. In particular, Biosynthetic Gene Cluster (BGCs) that code for specialized metabolites has gained significant interest due to their major role in modern society as a source of pharmaceutical drugs (e.g. antibiotics) and crop protection chemicals (1,2). These loci not only contain genes responsible for biosynthesis but often include auxiliary regions coding for regulatory and transporter proteins (2,3). Using signature genes and machine-learning-based methods, several computational frameworks have been developed to effectively detect hypothetical BGCs from genomic data, such as ClusterFinder, PRISM, DeepBGC, and antiSMASH (4–7). With these mature pipelines and the increase in publicly available genomes, a vast number of BGCs, both experimentally verified and hypothetical, have been catalogued in several databases. These include MIBiG, antiSMASH-DB, BiGFAM, ARTS-DB, and IMG-ABC (8–12). Unfortunately, much of this data remains unannotated. For instance, as little as 0.3% of the ~400,000 BGCs in IMG-ABC v5 are experimentally validated. Comparative genomic analysis can shed light on the functions of BGCs and their underlying genes. However, accessible online tools to allow scientists to perform custom comparative genomic analyses are lacking.

Gene cluster analysis methods for homology grouping, search, and visualisation are essential tasks to effectively leverage the available public resources. While tools such as BIG-SCAPE, BiGSLiCE, MultiGeneBlast and cblaster aid in gene cluster analysis, these demand local computational resources or require command-line experience (13–16). Due to the technological barrier, there is a need for a user-friendly and accessible platform for performing these analyses. Additionally, downstream methods for interpreting these results are often required. Visualisation and comparative genomic tools such as clinker and CORASON are capable of highlighting synteny or evolutionary relationships between BGCs; however, these also require expertise to operate and are not easily connected to homology search results (13, 17). To remedy this problem and provide an accessible, “BLAST-like” web server for gene clusters, we present CAGECAT (the CompArative GEne Cluster Analysis Toolbox).

The CAGECAT web server enables researchers to execute a full gene cluster analysis pipeline using customizable BLAST searches on up-to-date genomic databases. The service provides seamless connections between the search and visualisation modules, enabling execution, inspection, and fine-tuning of relevant search results. While some multi-gene search portals exist, such as ClusterScout and antiSMASH-DB, these only provide for model-based searching (e.g. Pfam) on predefined genome datasets, which often lag behind rapidly growing public genomic databases (9, 18). In addition to providing more up-to-date results, leveraging BLAST homology allows for refined control compared with model searches (e.g. identity and coverage), which can lead to more specific matches that aid in annotation, taxonomic distribution, or gene cluster evolution. Furthermore, with the interconnection of modules a user can accelerate result curation and downstream analysis, e.g. using gene neighbourhood estimation output to adjust intergenic distance thresholds to obtain more relevant matches. To our knowledge, we present the first free and publicly available web server for accelerated curation of homologous gene clusters with integrated downstream interpretation. By broadening accessibility of gene cluster analysis methods we hope this will lead to accelerated analysis and annotation of BGCs and contribute to the general knowledge of their subsequent products.

## IMPLEMENTATION AND AVAILABLE TOOLS

The aim of CAGECAT is to provide a platform to seamlessly connect gene cluster analysis tools in an accessible web server for search and interpretation of results. The search module leverages the cblaster pipeline, which utilises remote BLAST searches via NCBI’s servers as well as accelerated local Hidden Markov Model (HMM) based searches. Besides rapid similarity searches of entire BGC regions, cblaster provides several functions for gene neighbourhood estimation (GNE), sequence extraction, and visualisation (see Gilchrist et al. for a detailed description of methods) (16). The clinker pipeline is currently used for the visualisation module, which provides automated cluster alignment and homology annotations. CAGECAT has been designed to provide rapid interoperability between these functions, where homologous clusters of interest can be selected to be used in subsequent analysis. A graphical summary of tool interoperability is given in Figure 1.

**Figure 1:**
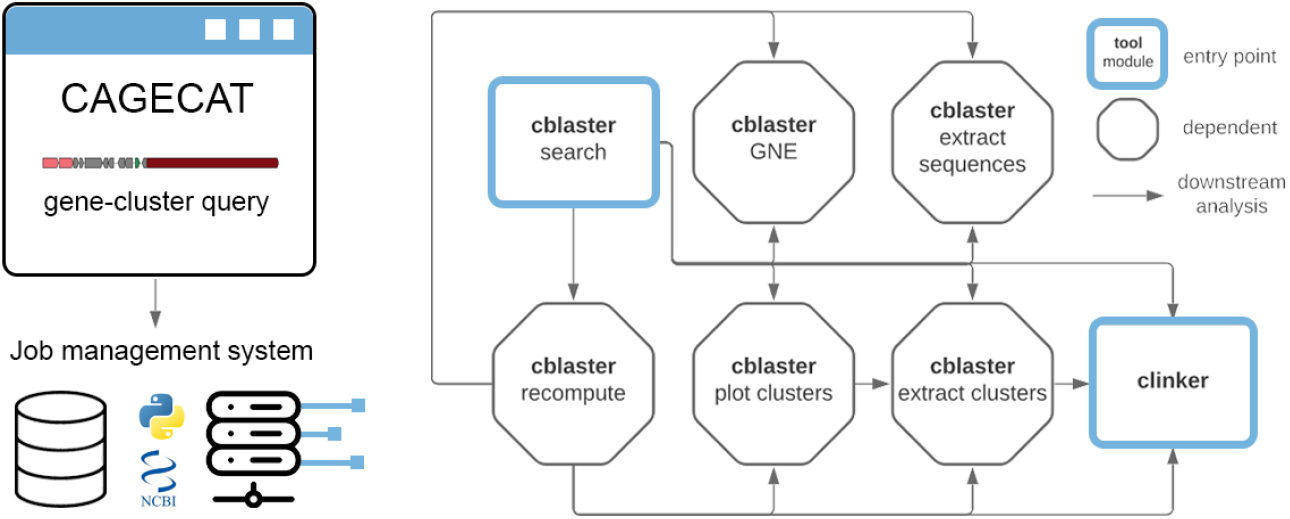
interoperability scheme of implemented functionality on CAGECAT. Blue outlined rectangles indicate entry points. Arrows indicate available downstream analyses from a module. Currently, a cblaster search/recompute job can be used for every downstream module, excluding a recompute job from being recomputed again. The clinker tool has no downstream analyses. For example, a possible workflow could be: cblaster search to cblaster recompute to cblaster plot clusters to selective clinker visualisation. This allows for fine-grained control of relevant matches for final visualisation and greatly improves user processing time.

### Databases for hidden markov model (HMM) searches

Searches for homologous gene clusters based on HMM profiles using cblaster require cblaster-generated HMM databases. Genus-specific Pfam databases were generated as detailed in supplemental methods resulting in 70 genera with 10 or more genomes for fungi, and 43 genera with 50 or more genomes. A custom script to fetch representative and reference genomes of prokaryotes and fungi was made using NCBI’s *e-search* utilities (19). To maintain CAGECAT’s free accessibility and storage, researchers will be required to use the command line version of cblaster or a local installation of CAGECAT to utilise custom HMM databases.

### Job Management

CAGECAT manages job submissions through a queue submission system, which processes jobs in a parallelizable first-in-first-out manner. Remote BLASTp queries are submitted to the NCBI API which leverages a scalable infrastructure allowing for multiple simultaneous searches (~10 requests/sec with an API key). By default, up to 15 jobs can be run in parallel to ensure stability and throughput. Upon job execution, the job command is constructed with the user-defined values of the input parameters and the appropriate pipelines are executed via Python. All output files are then stored and saved using a uniquely generated job ID. See supplemental methods for further technical details.

## RESULTS AND USER INTERFACE

### Input and Output

Two entry points for queries are currently implemented in CAGECAT for either gene cluster search via cblaster (search module) or visualisation via clinker (visualisation module) (Table 1). Input and output for other implemented modules are shown in Supplementary Table S1.

**Table 1:** current entry points of CAGECAT and their inputs and outputs. cblaster enables gene cluster searches and clinker creates publication-ready gene cluster visualisations. Additional downstream functions can be executed directly form results of previous session.

| Tool | Purpose | Input | Output |
| --- | --- | --- | --- |
| cblaster<br>(search module) | Search homologous gene clusters | - FASTA (protein)<br>- GenBank<br>- NCBI accessions<br>- HMM profile identifiers | -Interactive hit visualisation (HTML)<br>- Summary table<br>- Session file (JSON) |
| clinker<br>(visualisation module) | Visualise genomes containing homologous gene clusters (all vs. all) | - GenBank | -Interactive cluster visualisation (HTML) |
| recompute | Re-filter a previous search with new parameters | Session file output of cblaster search (JSON format) | -Interactive hit visualisation (HTML)<br>- Summary table<br>- Session file (JSON) |
| extract sequences | Extract sequences which contain a certain query |  | - Protein sequences (FASTA)<br>- Protein headers (TXT) |
| extract clusters | Extract selected clusters |  | - GenBank |
| Gene Neighborhood Estimation (GNE) | Check intergenic gap from results to optimize parameter |  | - Summary file (TXT) -<br>Interactive visualisation (HTML) |
| Plot clusters | Visualise a search session (align queries to clusters) |  | - Interactive visualisation (HTML) |

The search module allows for local files in either GenBank or FASTA format (protein sequences) to be uploaded and processed by the cblaster pipeline. Additionally, NCBI accession numbers can be used to submit a search query on the NCBI database, which can be combined with local searches using HMM profiles in predefined databases on CAGECAT. The input page (Figure S1) also contains optional parameters for selection of remote databases, search behaviour, and clustering of results. For the visualisation module, users can upload several genbank files or directly use outputs from the search module.

After completion of remote NCBI searches, users are presented with a cluster heatmap, which displays the absence/presence of each query protein sequence across the genomic hits (Figure 2A). As in the original cblaster, the results are sorted and colored based on BLAST similarity and number of matching proteins to the query cluster for rapid identification and comparison of homologous gene clusters across genomes. For the visualisation module, clinker will generate interactive gene cluster comparison figures with links drawn between similar genes on neighbouring clusters and shaded based on sequence identity (Figure 2B). Further details of these modules can be found at https://cagecat.bioinformatics.nl/tools/explanation and several example case studies for the cblaster output can be found in Gilchrist et al.

**Figure 2:**
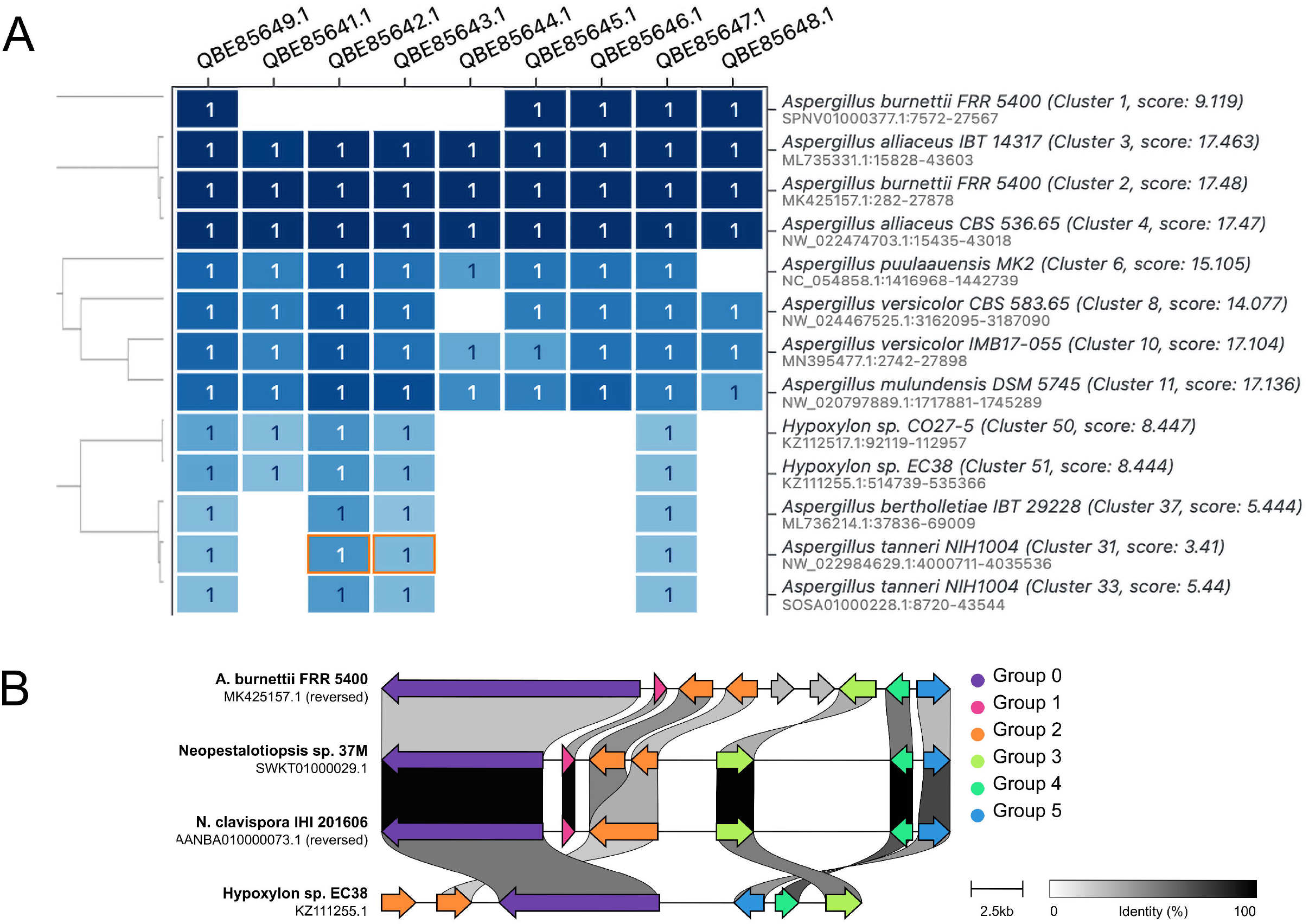
example output of CAGECAT’s entry point. Both modules create an interactive HTML visualisation which is displayed on each output page. (A) cblaster search: hit clusters are shown in a dendrogram (based on identity to query sequences). A darker tint of blue resembles a higher percentage identity of the query in the output cluster; (B) clinker visualisation: genes within a gene cluster are color-coordinated. Similar genes found in multiple clusters have links drawn between and are shaded based on sequence identity.

### Features and Interoperability

Users can download job results to their local computer within 30 days and output HTML files are displayed in-browser allowing for interactive inspection of results. The search module output allows for manual gene cluster selection to further curate results, which can be directly exported as genbank sequences. To accelerate analysis, CAGECAT provides interoperation between results and the available modules. Selections of output from the search module can be directly used as input for downstream analysis (e.g. to selectively visualise some results) or to recompute a search using different parameters (Figure 3). Notably, when genomic regions from the search module are used for analysis in the visualisation module, it will include all genes present within each genomic region that were not specified in the search query.

**Figure 3:**
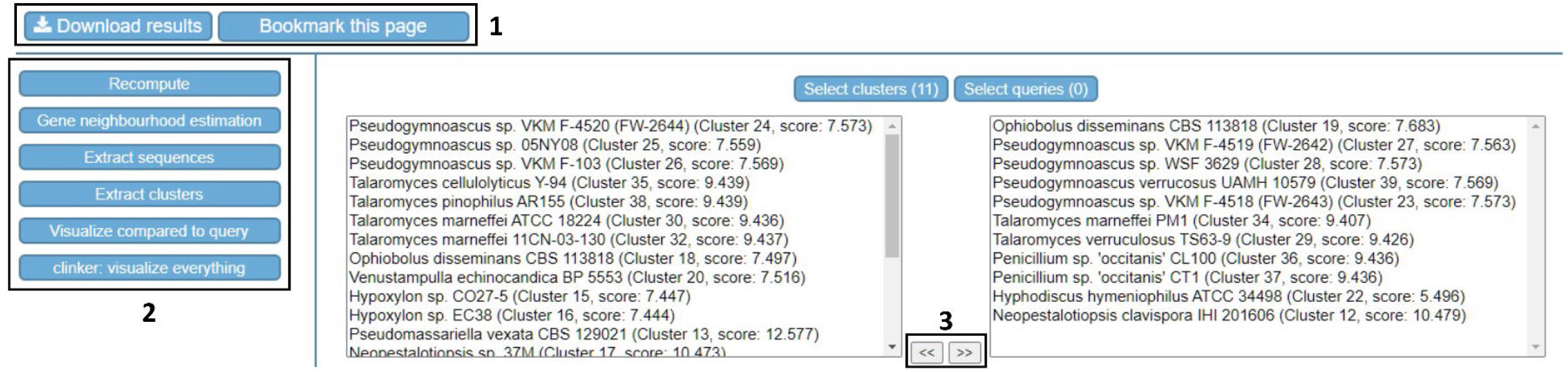
post-job execution screen for selective downstream analysis. 1: buttons to download results and save the current webpage to the browsers bookmark.; 2: available downstream analyses for the current analysis. Selected clusters and/or queries are temporarily saved when navigating to a downstream module; 3: manual selection of clusters for downstream analyses. Clusters/queries can be selected by moving them to the selected field using shown buttons. Available for cblaster search, recompute and plot clusters modules.

### Runtime and scalability

Remote search times are largely dependent on NCBI services which cannot be definitively benchmarked due to dependency on service traffic. However, processing of 346 queries over the 5-month user testing period showed an average search completion time under 8 minutes. Other functions such as clinker visualisation, recompute, gene cluster neighbourhood estimation, and cluster extraction all showed negligible processing time under 30 seconds (Supplementary Table S2)

## CONCLUSIONS AND FUTURE DIRECTIONS

With CAGECAT, we aim to lower the technical barrier to execute gene cluster analysis. Downstream analyses can be rapidly performed using the results of a previously executed job, which accelerates curation and comparative visualization. This service enables a quick search of whole gene cluster sequences against NCBI non-redundant or RefSeq databases that can be confined to a selected genus. Currently, two entry points exist to start analysing on CAGECAT: (I) finding homologous gene clusters using a query cluster and the cblaster search module, and (II) a visualisation of gene clusters using a set of query clusters and the clinker module. CAGECAT does not impact or interfere with the analysis capabilities of the implemented tools and acts as a bridge to allow for rapid retrieval of homologous gene clusters from continually updated public databases. We foresee CAGECAT being used by a wide audience to easily uncover homologous BGCs and provide publication-quality visualisations without the need for computational resources or programming expertise. Furthermore, CAGECAT is also useful for comparative analysis and discovery of gene clusters beyond those that encode the production of specialized metabolites, such as xenobiotic degradation pathways (20). With this web server, we aim to accelerate comparative analysis of gene clusters and provide an easy-to-use interface to help uncover clues for further study of BGCs encoding useful specialized metabolites as well as a starting point for investigating gene cluster evolution.

## AVAILABILITY

**Project name:** Comparative Gene Cluster Analysis Toolbox (CAGECAT)

**Project home page: https://cagecat.bioinformatics.nl**

**Operating system(s):** Linux / Platform independent via Docker

**Programming language:** Python

**Other requirements:** Python 3.8, Docker

**License:** MIT

**Source code: https://github.com/malanjary-wur/CAGECAT**

## LIST OF ABBREVIATIONS

API: Application programming interface
BGC: Biosynthetic Gene Cluster
CAGECAT: Comparative Gene Cluster Analysis Toolbox
CORASON: Core Analysis of Syntenic Orthologs to prioritize Natural Product BGCs
HMM: Hidden Markov Model
IMG-ABC: Integrated Microbial Genomes – Atlas of Biosynthetic Gene Clusters
MIBiG: Minimum Information about a Biosynthetic Gene cluster
NCBI: National Center for Biotechnology Information

## DECLARATIONS

### Authors’ contributions

M.B. developed and maintained web and core python architecture for CAGECAT. C.L.M.G provided cblaster / clinker integration support and product testing. Y-H.C and T.J.B contributed to testing and manuscript preparation. M.H.M and M.A. supervised and coordinated project development. All authors read and approved the final manuscript.

## Acknowledgements

We thank all researchers involved in beta testing from within the Bioinformatics group, Wageningen University, School of Molecular Sciences, The University of Western Australia

## Ethics approval and consent to participate

Not applicable

## Consent for publication

Not applicable

## Availability of data and materials

All data and materials are freely available via the updated git repository: https://github.com/malanjary-wur/CAGECAT as well as the release version used in this manuscript: https://github.com/malanjary-wur/CAGECAT/releases

## REFERENCES

1. Laich, F., Fierro, F., Cardoza, R.E. and Martin, J.F. (1999) Organization of the gene cluster for biosynthesis of penicillin in Penicillium nalgiovense and antibiotic production in cured dry sausages. Appl. Environ. Microbiol., 65, 1236–1240.

2. Medema, M.H. and Fischbach, M.A. (2015) Computational approaches to natural product discovery. Nat. Chem. Biol., 11, 639–648.

3. Crits-Christoph, A., Bhattacharya, N., Olm, M.R., Song, Y.S. and Banfield, J.F. (2020) Transporter genes in biosynthetic gene clusters predict metabolite characteristics and siderophore activity. Genome Res., 10.1101/gr.268169.120.

4. Cimermancic, P., Medema, M.H., Claesen, J., Kurita, K., Wieland Brown, L.C., Mavrommatis, K., Pati, A., Godfrey, P.A., Koehrsen, M., Clardy, J., et al. (2014) Insights into secondary metabolism from a global analysis of prokaryotic biosynthetic gene clusters. Cell, 158, 412–421.

5. Skinnider, M.A., Merwin, N.J., Johnston, C.W. and Magarvey, N.A. (2017) PRISM 3: expanded prediction of natural product chemical structures from microbial genomes. Nucleic Acids Res., 45, W49–W54.

6. Hannigan, G.D., Prihoda, D., Palicka, A., Soukup, J., Klempir, O., Rampula, L., Durcak, J., Wurst, M., Kotowski, J., Chang, D., et al. (2019) A deep learning genome-mining strategy for biosynthetic gene cluster prediction. Nucleic Acids Res., 47, e110.

7. Blin, K., Shaw, S., Steinke, K., Villebro, R., Ziemert, N., Lee, S.Y., Medema, M.H. and Weber, T. (2019) antiSMASH 5.0: updates to the secondary metabolite genome mining pipeline. Nucleic Acids Res., 47, W81–W87.

8. Kautsar, S.A., Blin, K., Shaw, S., Navarro-Muñoz, J.C., Terlouw, B.R., van der Hooft, J.J.J., van Santen, J.A., Tracanna, V., Suarez Duran, H.G., Pascal Andreu, V., et al. (2020) MIBiG 2.0: a repository for biosynthetic gene clusters of known function. Nucleic Acids Res., 48, D454–D458.

9. Blin, K., Shaw, S., Kautsar, S.A., Medema, M.H. and Weber, T. (2021) The antiSMASH database version 3: increased taxonomic coverage and new query features for modular enzymes. Nucleic Acids Res., 49, D639–D643.

10. Kautsar, S.A., Blin, K., Shaw, S., Weber, T. and Medema, M.H. (2021) BiG-FAM: the biosynthetic gene cluster families database. Nucleic Acids Res., 49, D490–D497.

11. Mungan, M.D., Blin, K. and Ziemert, N. (2022) ARTS-DB: a database for antibiotic resistant targets. Nucleic Acids Res., 50, D736–D740.

12. Palaniappan, K., Chen, I.-M.A., Chu, K., Ratner, A., Seshadri, R., Kyrpides, N.C., Ivanova, N.N. and Mouncey, N.J. (2020) IMG-ABC v.5.0: an update to the IMG/Atlas of Biosynthetic Gene Clusters Knowledgebase. Nucleic Acids Res., 48, D422–D430.

13. Navarro-Muñoz, J.C., Selem-Mojica, N., Mullowney, M.W., Kautsar, S.A., Tryon, J.H., Parkinson, E.I., De Los Santos, E.L.C., Yeong, M., Cruz-Morales, P., Abubucker, S., et al. (2020) A computational framework to explore large-scale biosynthetic diversity. Nat. Chem. Biol., 16, 60–68.

14. Kautsar, S.A., van der Hooft, J.J.J., de Ridder, D. and Medema, M.H. (2021) BiG-SLiCE: A highly scalable tool maps the diversity of 1.2 million biosynthetic gene clusters. Gigascience, 10.

15. Medema, M.H., Takano, E. and Breitling, R. (2013) Detecting sequence homology at the gene cluster level with MultiGeneBlast. Mol. Biol. Evol., 30, 1218–1223.

16. Gilchrist, C.L.M., Booth, T.J., van Wersch, B., van Grieken, L., Medema, M.H. and Chooi, Y.-H. (2021) cblaster: a remote search tool for rapid identification and visualization of homologous gene clusters. Bioinformatics Advances, 1.

17. Gilchrist, C.L.M. and Chooi, Y.-H. (2021) Clinker & clustermap.js: Automatic generation of gene cluster comparison figures. Bioinformatics, 10.1093/bioinformatics/btab007.

18. Hadjithomas, M., Chen, I.-M.A., Chu, K., Huang, J., Ratner, A., Palaniappan, K., Andersen, E., Markowitz, V., Kyrpides, N.C. and Ivanova, N.N. (2017) IMG-ABC: new features for bacterial secondary metabolism analysis and targeted biosynthetic gene cluster discovery in thousands of microbial genomes. Nucleic Acids Res., 45, D560–D565.

19. Entrez Programming Utilities Help [Internet]. Bethesda (MD): National Center for Biotechnology Information (US); 2010-.

20. Wisecaver, J. H., & Rokas, A. (2015). Fungal metabolic gene clusters—caravans traveling across genomes and environments. In Frontiers in Microbiology (Vol. 6).

