## Supplementary Table for "CAGECAT: The CompArative GEne Cluster Analysis Toolbox for rapid search and visualisation of homologous gene clusters"

### SUPPLEMENTAL METHODS

#### Back-end Server

CAGECAT runs within a Docker container and spawns multiple processes. User requests are passed via NGINX, a uWSGI server to FLASK which returns the appropriate web page. Submitted jobs are handled by FLASK by adding the job to the queue, created by *Redis*. The queued jobs are consumed and executed by the worker processes of the *rq* library which consumes these jobs in a first-in-first-out-manner. During job consumption, all custom values for parameters submitted by the user are integrated in the forged command, which is executed to perform the submitted analysis. Job information (i.e. title, type of job, status) are stored in a SQL database and queried from FLASK and the worker processes to aid in interoperable jobs.

HTML output is shown at the results page of a job if applicable and all output of a job is available for download. Executing a fine-tuned downstream analysis includes manual selection of hit clusters, which are sent along to the parameter page of the downstream analysis. Upon job submission, manually selected clusters are sent to CAGECAT modules to be used during analysis.

#### SERVER AND TECHNICAL SPECIFICATIONS

CAGECAT runs within an Ubuntu-based Docker (https://www.docker.com/) container on a Linux server (Ubuntu 18.04) equipped with two Intel® Xeon® Gold 6242@2.80GHz CPUs with 32 logical cores each and 768GB of RAM. The Docker container inherits all resources of the host. The *supervisord* library was used to supervise processes within the container, including *Redis* (job queue manager) and worker processes of the *rq* package (to consume jobs in queue). CAGECAT was written in Python using the web application framework Flask (https://flask.palletsprojects.com/) combined with dynamic web page rendering using Jinja2 server-side.
Job parameters are restricted to predefined options client-side and validated server-side to prevent execution of malicious code. Submitted jobs are stored in the client's cookies to navigate to results of previous jobs easily. These cookies do in no way infringe the user's privacy, as their only purpose is to facilitate easy navigation to previous jobs, and no other data besides the job type, id, title and submission date is stored.

#### HMM database creation

**NCBI e-search query:**

```
(((*[orgn] AND ("representative genome"[refseq category] OR "reference
genome"[refseq category])) AND (latest[filter] AND all[filter] NOT
anomalous[filter])))
```

Where * has been replaced with fungi and prokaryota, respectively.

After retrieval of all genomes the *makedb* command in cblaster was used to generate each set and these were associated with the corresponding genus entry in CAGECAT.

### SUPPLEMENTAL TABLES

Supplementary Table S1

| Function | Average time (min:secs) | Number of jobs |
|---|---|---|
| cblaster search | 07:20 | 346 |
| cblaster recompute | 00:05 | 65 |
| cblaster extract clusters | 00:24 | 66 |
| cblaster GNE | 00:06 | 11 |
| cblaster plot clusters | 00:01 | 40 |
| clinker | 00:14 | 118 |

### SUPPLEMENTAL FIGURES

**S1**

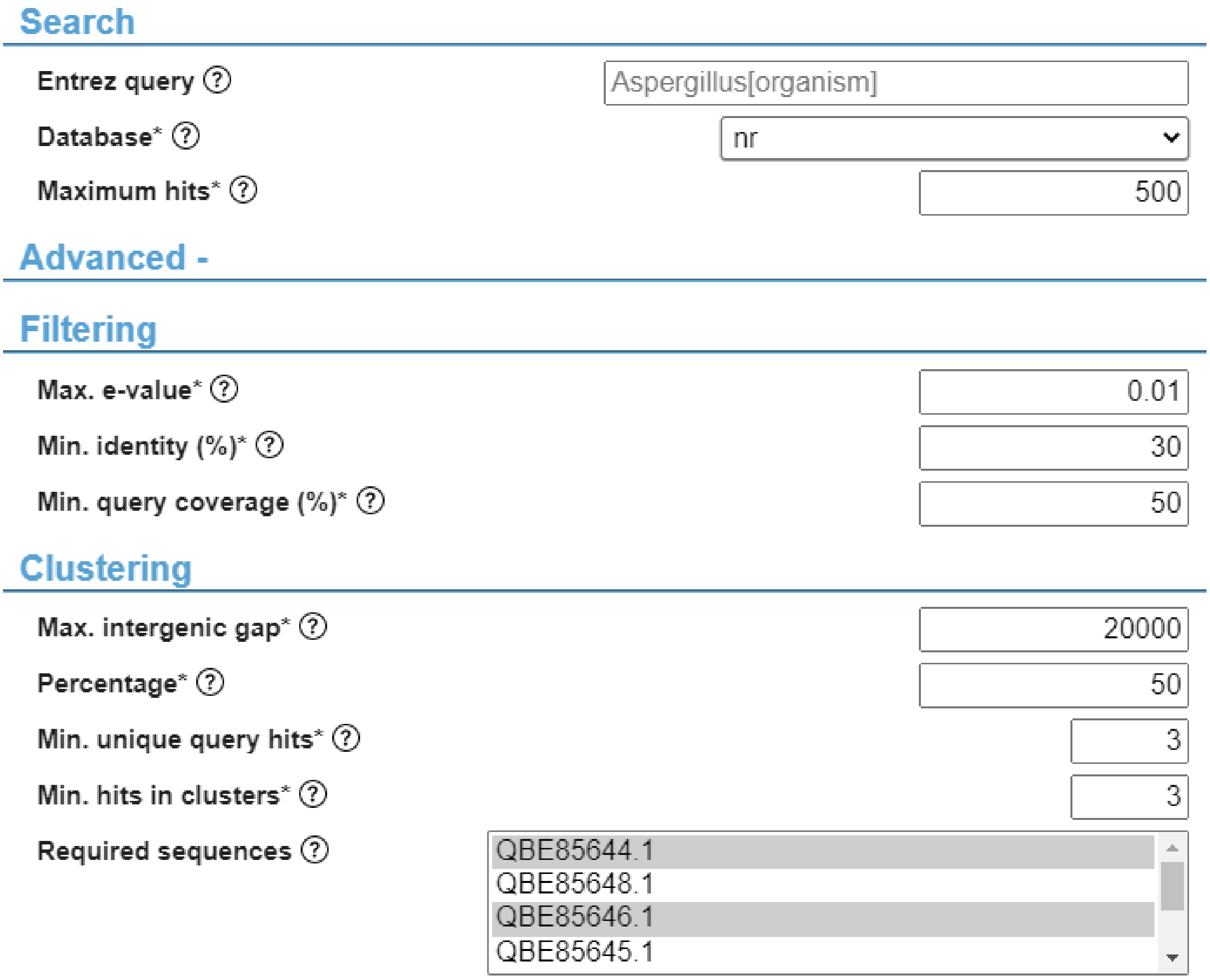


### TABLE AND FIGURES LEGENDS

**Supplemental figure S1: fields to set custom values for parameters of a cblaster search for gene clusters.** The advanced section shows many customizable parameters including a selection of query sequences that are obligated to be present in output clusters. Parameters in the advanced section will use default values (shown) if left untouched. Parameters with an * are obligatory and help texts are available per parameter by clicking the corresponding help button.

**Supplementary table S1: implemented cblaster modules on CAGECAT and their in- and output.** Shown cblaster modules use a session file created by either a cblaster search or recompute job, which contains all results of the gene cluster search and is used during analysis accordingly.

**Supplementary table S2: average execution time per module submitted to CAGECAT.** Homologous gene cluster searches using cblaster use <8 minutes and mostly consist of waiting on search results from NCBI.
